# Buzzed but not elated? Effect of ethanol on cognitive judgement bias in honeybees

**DOI:** 10.1101/2025.06.05.658067

**Authors:** Marta Golańska, Weronika Antoł, Magdalena Witek, Krzysztof Miler

**Affiliations:** Faculty of Psychology, University of Warsaw, Warsaw, Poland; Institute of Systematics and Evolution of Animals, Polish Academy of Sciences, Kraków, Poland; Museum and Institute of Zoology, Polish Academy of Sciences, Warsaw, Poland

**Keywords:** ethanol, cognitive bias, affective state, decision-making, learning, *Apis mellifera*

## Abstract

Ethanol is a neuroactive compound known to alter cognition, behavior, and physiology in a diverse range of vertebrate and invertebrate animals. It is also naturally present in many environments and might exert ecology-shaping influence on various organisms, such as honeybees feeding on fermenting nectar. In this study, we tested whether exposure of honeybees to low ethanol concentrations (1%) induces ‘pessimistic’ or ‘optimistic’ cognitive judgment bias (CJB), an information processing pattern thought to be related to affective processes. Bees were trained in a classical olfactory conditioning paradigm and then presented with conditioned stimuli and ambiguous odor mixtures to assess changes in decision-making under uncertainty. We additionally measured locomotor performance and ethanol absorption via spectrophotometric analysis of hemolymph. Ethanol levels peaked rapidly and remained elevated throughout the testing period. Despite this, ethanol had no detectable effect on choices in the CJB assay or locomotor behavior. These findings suggest that acute exposure to low-dose ethanol does not induce affect-like changes in honeybees. Our study represents the first test of pharmacologically induced judgment bias in an invertebrate and contributes to understanding the resilience of insect cognition to natural neuroactive compounds. The results highlight the limits and potential of CJB paradigms for probing affect-like states across taxa and support the honeybee as a valuable model in comparative cognition and the behavioral ecology of neuroactive substance exposure.

## 1. Introduction

Ethanol is a neuroactive substance occurring naturally in many environments, and some species have evolved mechanisms to tolerate or utilize it, despite its toxicity (Bowland et al. 2025; Miler 2025). Insects, in particular, frequently encounter ethanol in fermenting plant material. For example, fruit flies (*Drosophila melanogaster*) are attracted to ethanol-rich substrates, using it as a food source and to maintain a competitive advantage over less tolerant species (Merçot et al. 1994). The oriental hornet (*Vespa orientalis*) shows exceptional ethanol tolerance, possibly a result of coevolution with yeasts (Bouchebti et al. 2024). Ambrosia beetles, known for their advanced sociality and symbiosis with fungi, are attracted by ethanol emissions and select the colony sites based on ethanol concentration in the sap of candidate trees (Cavaletto et al. 2021). In addition, the compound has been found to promote the growth of the beetle’s nutritional symbiont (Lehenberger et al. 2021). These examples suggest that ethanol exposure can exert strong selective pressures and may have shaped behavioral and physiological adaptations across diverse insect taxa. Honeybees (*Apis mellifera*) also encounter ethanol in their environment, and it plays a significant role in their ecology (Abramson et al. 2025). Yeasts contribute to the fermentation processes involved in the bee bread production, a vital larval food source (Gilliam 1979; Detry et al. 2020 but see Anderson et al. 2014). Ethanol can also be present in flower nectar collected by foragers (e.g. Jakubska et al., 2005; Kevan et al., 1988; Schaeffer et al., 2017). Ethyl oleate, a pheromone produced from ethanol and oleic acid, regulates the division of labor in honeybee colonies (Castillo et al. 2012). Ethanol tolerance varies across worker castes, with higher resistance and alcohol dehydrogenase levels in foragers compared to workers staying inside the nest (Miler et al. 2021).

Much of what we know about the effect of ethanol on bee physiology, behavior, and cognition comes from studies that focus on honeybees as a potential model informing human alcohol addiction research. Honeybees have been found to prefer low ethanol concentrations in sucrose solutions (Abramson et al. 2000; Mustard et al. 2019) and show no conditioned taste aversion to ethanol (Varnon et al. 2018). Chronic ethanol consumption produces tolerance (Miler et al., 2018; but see Stephenson et al., 2021), and its discontinuation results in withdrawal-like symptoms in honeybees (Ostap-Chec et al., 2021; Yang, Z et al., 2025). Ethanol intake in honeybees can be reduced with disulfiram, a drug used in the treatment of alcohol dependence in humans (Abramson et al. 2003). Ethanol impairs honeybee motor function (Abramson et al. 2000; Maze et al. 2006; Ahmed et al. 2022) and learning performance (Abramson et al. 2000; Mustard et al. 2008). Social behaviors like trophallaxis or waggle and tremble dances are also affected (Bozic et al., 2006; Mixson et al., 2010). In addition, ethanol has been found to increase aggression (Wright et al. 2012; Giannoni-Guzmán et al. 2014), although not all studies support this finding (Abramson et al. 2000). Importantly, most studies use high ethanol concentrations (typically 2.5-10%), limiting their ecological relevance, as ethanol levels naturally encountered by honeybees are likely much lower (Wiens et al. 2008; Jones and Agrawal 2022).

Another limitation of the existing body of work is its limited scope in terms of the broadness of the cognitive traits investigated, focusing mostly on classical conditioning and memory. In recent years, more and more studies have uncovered the surprising cognitive complexity of honeybees, such as contextual and social learning, including also abilities that were thought to be unique to vertebrates, for example, metacognition, numerical cognition, conceptual learning, and attention-like processes (Collett et al. 1997; Leadbeater and Chittka 2007; Pahl et al. 2013; Avarguès-Weber and Giurfa 2013; Perry et al. 2017). One particularly interesting phenomenon is cognitive judgment bias (CJB), which measures decision-making under uncertainty. The CJB paradigm offers a promising tool for investigating affect-like processes as well as their evolutionary basis and function. Originally developed for rodents (Harding et al. 2004), CJB tests assess whether animals show ‘optimistic’ or ‘pessimistic’ biases when interpreting ambiguous cues. It has been adapted for invertebrates, including honeybees (Bateson et al. 2011). In these tests, animals are conditioned to associate two stimuli with a rewarding stimulus, like food, and a punishment, like an unpalatable substance, and are later presented with the learned stimuli and novel, ambiguous ones. Reactions to these ambiguous cues might resemble the animal’s responses to the rewarded or punished stimuli, suggesting ‘optimistic’ or ‘pessimistic’ biases, respectively. In honeybees, ‘pessimistic’ biases have been induced by stressors (shaking; Bateson et al. 2011; Schlüns et al. 2017), and ‘optimistic’ biases have been investigated in their relatives, bumblebees (Solvi et al. 2016; Strang and Muth 2023; Romero-González et al. 2025). Changes in CJB are commonly interpreted as indicators of affect or emotion-like changes, following the original formulation of the test (Harding et al. 2004; Mendl et al. 2009; Neville et al. 2020; Lagisz et al. 2020, but see alternative interpretation by Baracchi et al. 2017). Whether insects possess emotional states or their analogues remains highly debated (see Baracchi et al. 2017; Perry and Baciadonna 2017). However, CJB responses in invertebrates provide a behavioral window into valence-based processing. Following Mendl and Paul (2020), we adopt a broad definition of ‘affect’ that includes all valence-related processes (‘pleasantness’ or ‘unpleasantness’), including emotions, moods, and their possible manifestations (physiological or behavioral), whether or not they entail conscious experience. This allows for comparisons across taxa without necessarily implying that conscious experience must be involved.

Some pharmacological substances have been shown to impact CJB across several, mostly mammalian, species (see Lagisz et al. 2020, for review and meta-analysis). Ethanol is a potential pharmacological moderator in this context. It is known to introduce affective changes in humans, acting as euphoriant in lower, anxiogenic in higher doses, and increasing aggression in some individuals as well (Freed 1978; Chermack and Giancola 1997). Rodent studies largely find similar patterns, mostly using traditional behavioral assays (Van Erp and Miczek 1997; Valdez et al. 2002). Ethanol has never been tested as a mood-manipulating agent in a CJB test; however, one recent study linked optimistic-like responses to ethanol self-administration in rats (Cieslik-Starkiewicz et al. 2024). Moreover, no study has tested whether ethanol administration can alter CJB in invertebrates, including those that are exposed to it under natural conditions.

Because ethanol is a neuroactive substance, even species with relatively high ecological tolerance may still exhibit transient changes in decision-making or motivational processes upon exposure. Testing this provides insight into the robustness of insect cognition to naturally occurring neuroactive compounds. In the present study, we investigated whether a low concentration (1%) of ethanol can induce CJB in honeybees. We hypothesized that ethanol-exposed bees would exhibit bias, that is, respond differently to at least one ambiguous cue compared to controls, while maintaining normal responses to learned stimuli. Additionally, we assessed locomotor behavior and ethanol absorption via hemolymph sampling to rule out non-specific performance effects and confirm effective dosing.

## 2. Methods

### 2.1. Cognitive judgment bias (CJB) test

#### 2.1.1. Study species and collection procedure

Bees were collected from three families from the experimental apiary located in Kraków, Poland, over 16 days. Animals were collected in one or two blocks daily between 07:30 am and 12:25 pm, with each collection lasting up to 15 minutes. During the procedure, entrance to the hive was partially blocked by a foam barrier, allowing the experimenter to select and capture congregating bees. Only pollen-carrying individuals were chosen to ensure their forager status. Up to 30 bees were restrained per block. Bees were left for one hour to acclimate and strengthen feeding motivation. From each block, 12 bees were selected for the conditioning phase based on their overall condition, absence of proboscis or antennal defects, adequate restraint, and prompt proboscis extension upon brief antennal stimulation with sucrose solution. The remaining bees were released (see Supplementary Information 1 for a visualization of the exclusion process).

#### 2.1.2. Olfactory conditioning

Selected bees underwent olfactory conditioning using the proboscis extension response (PER) paradigm (Scheiner et al. 2013). This classical conditioning paradigm utilizes a reflexive extension of the proboscis in response to sucrose contact and is widely used for appetitive conditioning in honeybees (Giurfa and Sandoz 2012). Although PER is a reflex, it is known to be modulated by many factors, including motivational ones (Menzel, R 1990), and it has been successfully used in bee CJB protocols before (Bateson et al. 2011; Schlüns et al. 2017). Here, the standard appetitive conditioning protocol also included discrimination learning with NaCl as an aversive (de Brito Sanchez et al., 2015). One odor (2-octanone, >99%, Thermo Scientific) was paired with an appetitive unconditioned stimulus (0.5 μl drop of 1.5 M aqueous sucrose solution; hereafter US), while a second odor (1-hexanol, >98%, Tokyo Chemical Industry) was paired with an aversive unconditioned stimulus (0.5 μl drop of 0.5 M aqueous NaCl solution; US). Each odor was delivered from a 24 ml syringe containing a cotton pad soaked with 6 µl of the odorant. The end of the syringe was placed about 0.5 cm from the bee’s head, and air was expelled for 4 seconds. In the final second of odor delivery, US or US was presented, lasting 3 seconds in total (Matsumoto et al. 2012). The antennae were stimulated with the solution, and if PER was elicited, a drop was placed on the proboscis using a pipette.

Each bee underwent 16 conditioning trials (8 per odor) in a fixed sequence (CS –CS –CS –CS –CS – CS –CS –CS –CS –CS –CS –CS –CS –CS –CS –CS; as used by Schlüns and colleagues (2017), where CS and CS stand for the odors associated with the US (sugar solution) and US (salt solution), respectively. Bee responses during the first 3 seconds of odor presentation (before US delivery) were recorded as 1: full PER, 0: no PER, or partial extension of the proboscis. Inter-trial intervals were 6 minutes. Bees were included in the experiment if they displayed correct responses in at least four of the final six trials for each odor. Lack of PER to hexanol and full PER to octanone were considered correct responses.

#### 2.1.3. Group assignment and experimental manipulation

Bees that met the learning criteria were randomly assigned to either the control group (fed 10 µl of 1.5 M aqueous sucrose solution) or the ethanol group (fed 10 µl of 1.5 M aqueous sucrose solution with 1% ethanol). For each block, group sizes were balanced wherever possible, or when the number of bees was odd, the more numerous group was chosen randomly. The individuals were then randomly assigned to the groups. Successfully fed bees were left for one hour before testing to allow ethanol absorption and improve feeding motivation. Bees not meeting the learning criteria were marked to prevent reuse and released.

#### 2.1.4. Testing procedure

The successfully fed bees participated in the CJB test comprising five odor presentations (testing trials). Each odor was presented for four seconds, and PER responses were coded as 1: full PER and 0: lack of PER or partial response. No US were delivered, but a pipette was held nearby to mimic the conditioning context. The odors included the two conditioned odors (6:0 octanone:hexanol = positive; 0:6 = negative) and three mixtures: 4:2 (near-positive), 3:3 (neutral), and 2:4 (near-negative). Odors were delivered as in the conditioning phase. The experimenter was blind to the odor and the testing order, which was randomized per block.

Each testing trial was preceded by two reminder trials (US, US) to limit extinction of the conditioned responses (Sandoz and Pham-Delègue 2004). The sequence was RT –RT –**T1**-RT –RT –**T2**-RT –RT –**T3** –RT –RT –**T4**-RT –RT –**T5** (RT and RT = reminder trials with appetitive and aversive US paired with the conditioned odors; **T1-5** = testing trials). The full procedure lasted 44 minutes, with a 3-minute inter-trial interval. After testing, bees were marked and released, and odor identities were unblinded.

### 2.2. Locomotion test

Bees were collected from seven families from the same apiary over nine days (between 07:00 am and 09:30 am, in two blocks), with one bee collected per minute for up to 30 minutes. Only pollen-carrying individuals were selected, using the same method as described above, and they were restrained promptly after being caught. Bees were left for three hours to acclimate and approximate the conditioning phase duration in the CJB test. The first 15 and last 15 bees of each block were assigned to the ethanol-fed or control group, with group assignment counterbalanced across families.

Bees were fed 10 µl of either 1.5 M aqueous sucrose solution (control) or the same solution with 1% ethanol (ethanol-fed). Animals were excluded based on visible leg or mouthpart defects, poor condition, incomplete consumption, or mortality (for the full inclusion/exclusion process for the individual bees participating in the test, see Supplementary Information 1). Fifty minutes post-ingestion, bees were released from restraint and placed under a Petri dish (14.5 cm diameter, 1 cm high) for 10 minutes to acclimate. One hour after feeding, a 20-minute video of their locomotion inside the dish was recorded. Videos were processed using AnimalTA (Chiara and Kim 2023) to extract the total distance travelled, average speed while moving, and meander (average of the change in direction, or turning angle, divided by the distance travelled for each frame).

### 2.3. Ethanol levels in hemolymph

Bees were collected from four families from the same apiary over six days. The animals were collected for one hour, starting at 09:00 am, and assigned to one of four ethanol-fed groups, 15 individuals per group. Only pollen-carrying bees were selected (using the method described in previous sections), then restrained and fed 10 µl of 1.5 M sucrose solution containing 1% ethanol. Four ethanol-fed groups differed in the time of hemolymph extraction after feeding (30, 60, 120, or 240 minutes). An additional control group (N = 15) received a sucrose-only solution and was sampled 30 minutes post-ingestion.

All bees were restrained for two hours before feeding to acclimate and mimic the CJB test procedure. The animals that did not ingest the entire solution were not used (again, for the full inclusion/exclusion process for the individual bees participating in the test, see Supplementary Information 1). Hemolymph was extracted using the antennal method (Borsuk et al. 2017) and collected in end-to-end 10 µl microcapillaries, then frozen until analysis.

Samples were processed using the Ethanol Assay Kit, Megazyme, following the manufacturer’s protocol with some modifications. This method relies on spectrophotometric detection of NADH, a product of two enzymatic reactions involving ethanol as a substrate. For each reaction, we used the minimal volume of 3 µl of hemolymph from two samples pooled within a group, or, if the volume was sufficient, from a single individual. The absorbance at 340 nm was measured twice: before (A1) and 5 min after adding the enzyme initiating the reaction (A2). The difference A2-A1 was used to calculate ethanol concentration in the sample, based on the calibration curve with blank, performed with each reaction plate.

### 2.4. Statistical analysis

All analyses in our experiments were performed using *R* (R Core Team, 2014). Model diagnostics were conducted using the packages performance (Lüdecke et al. 2021) and DHARMA (Hartig, 2024), and the emmeans package was used to obtain marginal means (Lenth, R and Piaskowski, J 2025).

#### 2.4.1. CJB test

We analyzed PER responses (1 = full PER; 0 = absent/partial PER) using generalized mixed-effects binomial models. Odor stimuli consisted of discrete mixtures (0, 2, 3, 4, 6 parts octanone), and so we first compared a model assuming a strictly linear effect of octanone proportion with a model treating odor as a categorical predictor. This comparison indicated that the linear model fit worse, suggesting nonlinearity in the relationship between odor mixture and PER probability. We therefore used a generalized additive mixed model (GAMM; mgcv package; Wood, S. N., 2017) with a smooth term for odor mixture and a random-effect smooth for individual bee ID to account for repeated measures. The model included group (ethanol-treated vs. control) as a fixed effect and a common smooth for odor mixture. To test judgment bias, we compared this model to a model allowing group-specific smooths using ANOVA. Model-estimated marginal means and 95% confidence intervals at the tested odor mixtures were obtained from the fitted model for visualization and contrasts. A total of 736 observations from 148 bees were analyzed. Data for one individual was discarded due to experimenter error. We additionally explored models that included date, hour, odor presentation order, and family (as fixed effects), but these covariates were all non-significant and did not improve model fit and were excluded from consideration. As a robustness check, we fitted the GAMM described above on the data with a variant coding of PER (1 = full or partial PER, 0 = absent PER) in order to verify whether decisions regarding coding of the outcome variable meaningfully impact the result.

For analyzing learning trials, we considered responses from all 274 bees participating in the learning procedure, for a total of 4230 trials. In order to analyze conditioning acquisition, we used a binomial generalized linear mixed model with a logit link (package lme4; Bates, D et al., 2015). PER (1 – full PER, 0 – absent/partial PER) was modeled as a function of conditioned stimulus type (CS vs. CS), conditioning trial pair (1-8, treated as a categorical factor), and their interaction. Individual bee ID was included as a random intercept to account for repeated measurements. Model-based estimated marginal means and 95% confidence intervals were obtained.

#### 2.4.2. Locomotion test

We analyzed the effect of ethanol on three locomotor variables: distance travelled while moving, average speed while moving, and meander (change in direction per unit distance). Movement was defined using a threshold of 0.3 cm/s to minimize the influence of tracking imperfections, with perspective correction and default smoothing applied (see Chiara and Kim 2023 for explanations of the parameters). Data from four individuals were excluded due to poor physical condition. Average speed while moving was analyzed using a linear model (package nlme; Pinheiro, B, 2000) with treatment group (control vs. ethanol-fed) as a predictor. Distance travelled while moving was analyzed using a linear model fitted with generalized least squares on log-transformed distances with group-specific residual variance to account for heteroscedasticity. Meander was analysed using a linear model on log-transformed values. Model estimates were obtained with 95% confidence intervals. The final sample size was 280 individuals.

#### 2.4.3. Ethanol levels in hemolymph

Ethanol concentrations in hemolymph were analyzed using a linear model fitted to log-transformed values after applying a log(1 + x) transformation to accommodate zero values arising from measurements below the detection limit. Sampling group (control, 30-, 60-, 120-, or 240-minute post-ingestion) was included as a fixed effect. Family, assay plate, and the number of individuals pooled per sample were included as covariates to account for noise. Model-based estimated marginal means were computed for each group, and contrasts were used to compare ethanol-fed groups to the control as well as to assess differences among ethanol-fed time points. Model-based estimates were obtained with 95% confidence intervals. In total, 55 hemolymph samples from 105 individuals were analyzed.

## 3. Results

Responses in the cognitive judgment bias (CJB) test were strongly influenced by odor mixture composition (Figure 1). In both control and ethanol-fed bees, the probability of PER increased nonlinearly with increasing proportion of the rewarded odor (GAMM smooth terms: control: χ² = 65.53, p < 0.001, ethanol-fed: χ² = 84.95, p < 0.001). Ethanol treatment had no significant main effect on PER probability (estimate = −0.27 ± 0.42 SE, z = –0.66, p = 0.51). Importantly, ethanol did not alter the shape or slope of the odor-response function, as indicated by the absence of group-specific differences in the nonlinear odor smooths (χ² = 5.78, p = 0.22). Responses to ambiguous odor mixtures (near-positive, neutral, and near-negative cues) were indistinguishable between treatment groups, providing no evidence for ethanol-induced CJB. For the three ambiguous mixtures, the odds-ratio confidence intervals were 2:4 (OR = 0.90, 95% CI [0.28, 2.95]), 3:3 (OR = 0.98, 95% CI [0.39, 2.42]), and 4:2 (OR = 0.91, 95% CI [0.38, 2.19]), incompatible with meaningful ethanol-induced shifts in ambiguous-cue responding, although modest effects remained possible. No differences in learned odors suggested no post-ingestion changes in memory or proboscis mobility (see Supplementary Information 2 for all contrasts). The model explained 51.6% of the deviance in PER responses. No major differences in the model parameters with variant outcome variable coding were detected (see Supplementary Information 2).

**Figure 1.**
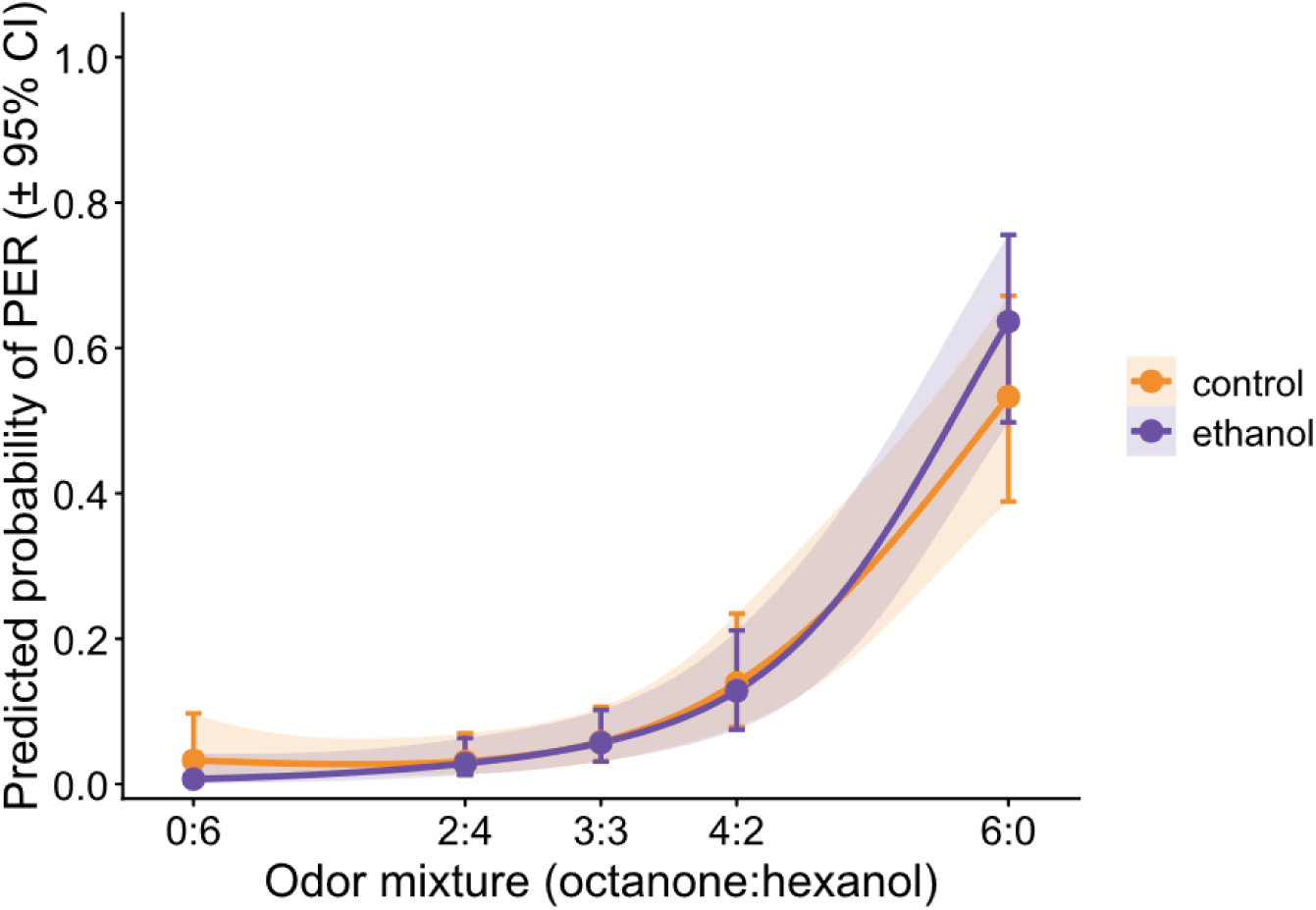
Predicted probability of PER across odor mixtures in the CJB test. Curves represent predictions from a binomial generalized additive mixed model, with shaded bands indicating 95% confidence intervals. The x-axis shows the proportion of octanone (appetitive CS) in the odor stimulus, including conditioned cues (6:0 and 0:6) and ambiguous mixtures (4:2, 3:3, 2:4). Both groups show similar nonlinear odor-response functions, with no evidence of ethanol-induced shifts in responses to ambiguous cues.

In the locomotor activity test, control and ethanol-fed bees showed similar performance across the measured parameters (Figure 2). Average speed while moving did not differ between groups (control: 3.30 cm/s [95% CI 3.13, 3.47]; ethanol-fed: 3.19 cm/s [95% CI 3.02, 3.36]; t = –0.88, p = 0.38). For distance travelled while moving, there was a non-significant tendency for ethanol-fed bees to travel less than controls (control: 3470 cm [95% CI 3193, 3772]; ethanol-fed: 3018 cm [95% CI 2654, 3432]; t = –1.80, p = 0.07). Meander while moving also did not differ between groups (control: 227 deg/cm [95% CI 214, 240]; ethanol-fed: 229 deg/cm [95% CI 216, 242]; t = 0.20, p = 0.84).

**Figure 2.**
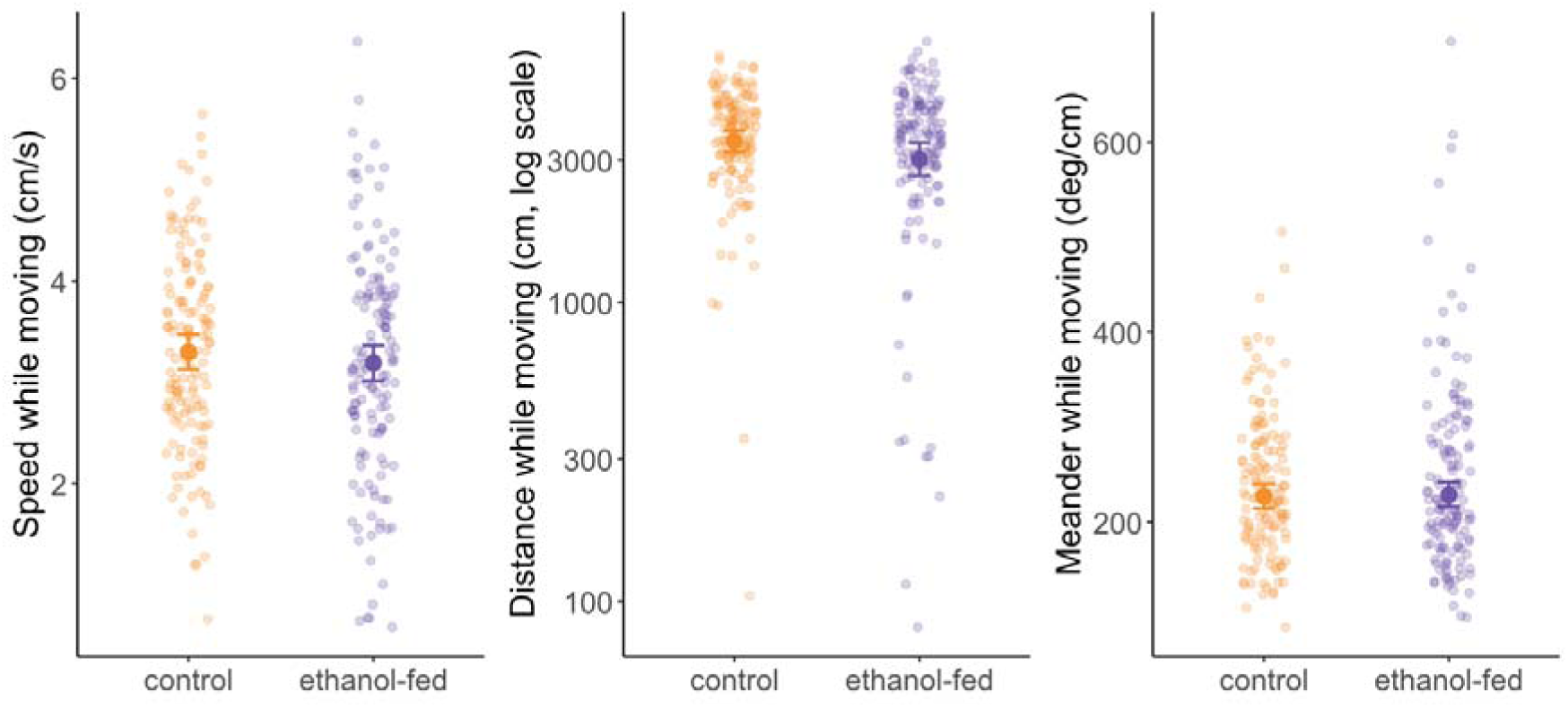
Locomotor activity in control and ethanol-fed honeybees. Raw individual values are shown as semi-transparent points; larger points and error bars indicate model-estimated marginal means ± 95% confidence intervals. Speed while moving was analyzed with a linear model. Distance travelled while moving was analyzed using generalized least squares on log-transformed distances with group-specific residual variance. Meander while moving was analyzed with a linear model on log-transformed values. Ethanol treatment did not affect speed, showed a non-significant tendency toward reduced distance, and did not affect meander.

Hemolymph ethanol concentrations differed among sampling groups as a function of time since ingestion (F_4,45_ = 13.59, p < 0.001; Figure 3). Compared to control bees, ethanol-fed ones showed significantly elevated ethanol concentrations at 30 minutes (estimate = 0.65, SE = 0.09, t = 6.98, p < 0.001), 60 minutes (estimate = 0.49, SE = 0.09, t = 5.24, p < 0.001), and 120 minutes (estimate = 0.40, SE = 0.11, t = 3.74, p = 0.005) post-ingestion. At 240 minutes after ingestion, ethanol concentration was only slightly elevated (estimate = 0.30, SE = 0.12, t = 2.59, p = 0.090). No additional effects of family, assay plate, or pooling of individuals were detected (see Supplementary Information 2 for full results and contrasts). Together, these results indicate that hemolymph ethanol levels remained elevated throughout the behavioral testing period (30-120 minutes) and declined toward baseline within four hours of ingestion.

**Figure 3.**
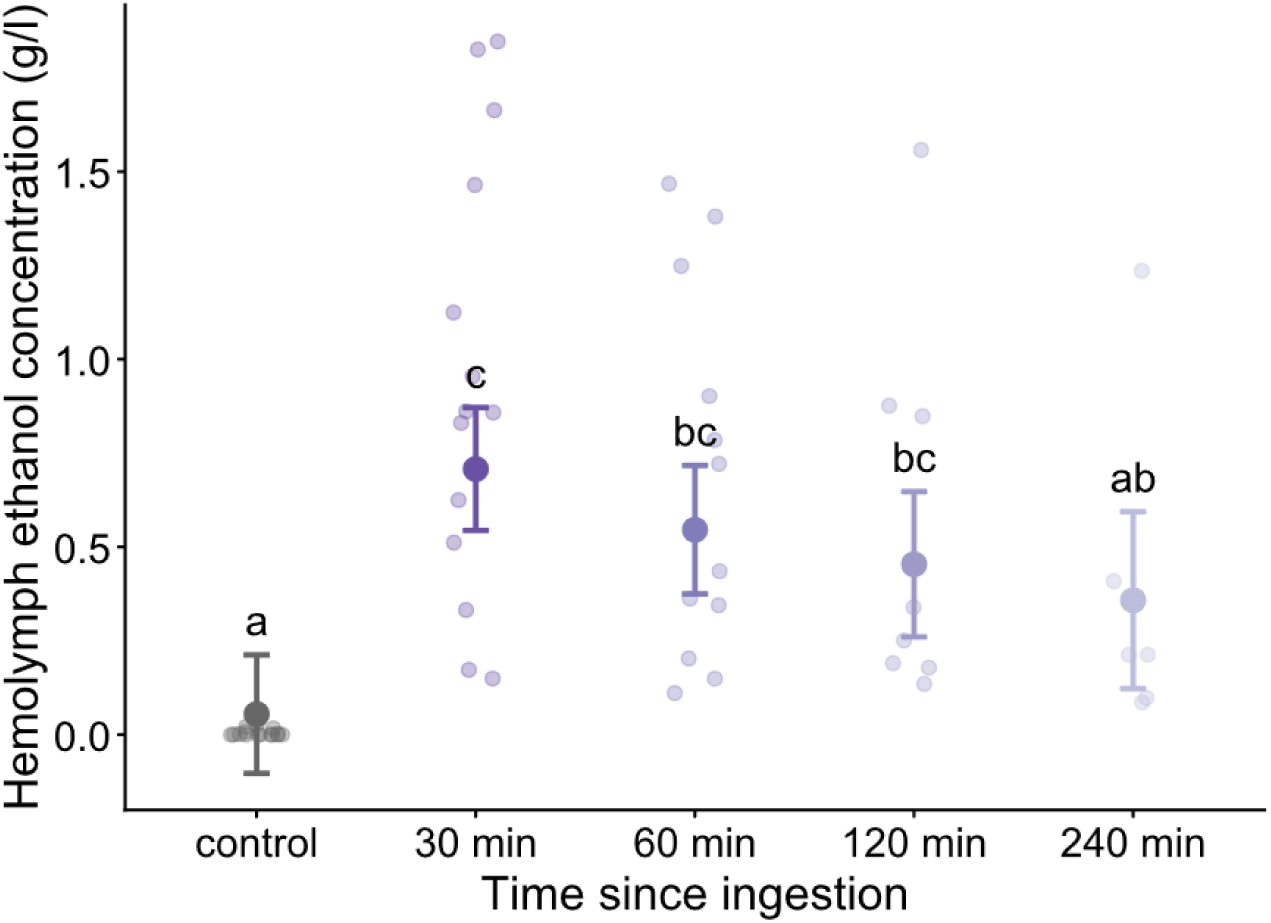
Hemolymph ethanol concentration following ingestion of ethanol-containing sucrose solution. Bees were sampled at 30, 60, 120, and 240 minutes after feeding on a 1% ethanol solution; control bees received sucrose without ethanol and were sampled 30 minutes after ingestion. Semi-transparent points represent individual hemolymph samples. Larger points and error bars indicate model-estimated marginal means ± 95% confidence intervals from a linear model fitted to log-transformed ethanol concentrations, adjusted for family, assay plate, and pooling of individuals. Different letters indicate significant differences among sampling groups based on Tukey-adjusted pairwise comparisons. Ethanol concentrations were elevated at 30-120 minutes post-ingestion and approached control levels by 240 minutes.

Analysis of responses during the conditioning phase confirmed effective acquisition. There was a significant main effect of conditioned stimulus type (CS vs. CS; χ² = 7.94, p = 0.0048), a strong effect of conditioning trial pair (χ² = 274.56, p < 0.001), and a pronounced CS type × trial interaction (χ² = 150.95, p < 0.001). PER probability increased sharply across successive trial pairs for the rewarded odor (CS), while responses to the punished odor (CS) remained low and diverged progressively from CS across training. This pattern indicates robust associative learning and reliable discrimination between conditioned stimuli across the conditioning sequence (see Supplementary Information 2 for predicted probabilities of PER across conditioning and Figure S1).

## 4. Discussion

Our study investigated whether ethanol exposure influences cognitive judgment bias (CJB) in honeybees. We found no evidence that ethanol affected bees’ responses to ambiguous cues, suggesting an absence of judgment bias. Additionally, we observed no detectable influence on bees’ responses to the conditioned stimuli themselves, and no clear impairment of general motor function. Ethanol ingestion and absorption were confirmed by spectrophotometric analysis of hemolymph samples, which showed elevated levels during the behavioral testing window. Together, these findings suggest that acute exposure to low doses of ethanol does not produce detectable changes in judgment bias within a PER-based CJB paradigm in honeybee.

Our findings contrast with earlier investigations using similar paradigms, which have demonstrated that invertebrates can show affect-like states. For instance, Schlüns et al (2017) found that honeybees exposed to a physical stressor (shaking) exhibit pessimistic biases (building on the pioneering, but less methodologically clear report by Bateson et al., 2011). CJB has also been described in bumblebees (Solvi et al. 2016; Strang and Muth 2023; Procenko et al. 2024; Romero-González et al. 2025) and fruit flies (Deakin et al. 2018). To our knowledge, however, this is the first study to investigate whether pharmacological intervention may induce CJB in an invertebrate. The absence of an effect is compatible with two major non-exclusive interpretations: that ethanol may not reliably modulate affect-like processing in bees at this dose and time scale, or that the PER-based CJB paradigm may be relatively insensitive to ethanol-induced state changes. We considered whether the absence of CJB could reflect insufficient ethanol absorption, but this is unlikely. As in earlier studies (Bozic et al. 2006; Maze et al. 2006), ethanol was absorbed rapidly and persisted in hemolymph during the behavioral testing period. Conditioning dynamics also indicated robust acquisition and discrimination, supporting the effectiveness of the learning procedure.

Dose and exposure regime are plausible explanations. Ethanol’s influence may not manifest at low exposure, even though 1% ethanol has been reported to affect honeybee behavior in some contexts (e.g. Bozic et al. 2006; Ahmed et al. 2022). Our own data suggests a trend for a change in gross locomotor behavior after 1% ethanol consumption. Higher concentrations might induce measurable bias, and further investigation is warranted, but such studies should also account for the known confounding effects of ethanol on learning and motor performance. From an ecological and apicultural perspective, exploring a wider ethanol range is relevant because although nectar in temperate climates is unlikely to reach high ethanol concentrations, bees may encounter high doses in particular environments, especially human-modified ones such as orchards where fermenting fruits are available (Ostap-Chec et al., 2025). It remains unclear how often and under what circumstances bees supplement their diet with fallen fruits (Abramson et al. 2025) and whether this exposure meaningfully affects their decision-making or welfare. Changes to affect and decision-making might also only be apparent after chronic exposure, instead of a single acute dose, as in our study. Another major possible explanation is that the binary measure of PER might not be sensitive enough to detect the change in behavior, and newer alternatives, such as quantifying the strength of the proboscis response (see Giurfa & Sandoz, 2012 for discussion), might yield different results. PER is a reflex, and its suitability for detecting subtle affect-related modulation may be generally questioned. Additionally, inter-individual variability may have obscured a small existing effect (Van Der Goot et al. 2021).

Our findings are consistent with the broader idea that honeybees may be relatively robust to low ethanol doses present in their environment. Natural ethanol production dates back to the late Cretaceous, when yeast gained the ability to ferment sugars in fleshy fruits of early angiosperms (Benner et al. 2002). Selection could plausibly favor physiological mechanisms that mitigate ethanol’s deleterious neural effects, particularly in foragers that are most exposed to the substance. In fact, they are known to be the most ethanol-resistant caste and possess elevated alcohol dehydrogenase levels (Miler et al. 2021). If bees have evolved physiological mechanisms that buffer ethanol’s neural impact, then affect-related cognitive processes might be comparatively resistant to low doses, which could help explain the absence of detectable bias in our paradigm despite sustained hemolymph ethanol levels. The persistence of ethanol in hemolymph is likely due to the gradual passage of food from its storage in the crop to the stomach, where it is digested (Crailsheim 1988). This makes low-dose robustness potentially consequential: even a single ethanol-containing meal could influence bees throughout a foraging bout, a vital but risky task that poses challenges to successful navigation, learning, and predator avoidance. However, despite ethanol’s potent, cross-species impact on neurotransmission (Phillips and Shent 1996), our results suggest that many low-dose behavioral consequences, if present, are hardly detectable.

Our findings help reconcile mixed reports in the literature. Some previous studies align with our findings, showing no effect of ethanol on bee learning (Abramson et al. 2000) or flight performance (Ostap-Chec et al. 2025). However, other works report behavioral changes even at 1% ethanol, including altered flight kinematics (Ahmed et al. 2022), delayed return to the hive, reduced trophallaxis and self-grooming, as well as changes in social communication (Bozic et al. 2006). One possibility is that low concentrations of ethanol impact aspects of body movement that bees can compensate for, making the effects undetectable at coarser scales (e.g. speed). Another possibility is that some consequences of ingesting low ethanol concentrations are only apparent on the group level and might reflect collective decision-making, for example, to reduce foraging when many individuals carry ethanol-containing food at the same time. For instance, Bozic and colleagues (2006) reported effects of colony-level inebriation when multiple foragers consumed ethanol simultaneously. Future studies should explore how low-dose ethanol concentrations influence social dynamics and collective behavior in honeybees.

## 5. Conclusion

This study investigated the impact of ethanol on cognitive judgment bias in honeybees. We show that acute exposure to 1% ethanol produces no detectable judgment bias in honeybees when CJB is assessed through binary PER responses to ambiguous odor mixtures and does not produce any major changes in gross locomotor performance. Ethanol presence was confirmed in the hemolymph for prolonged periods post-ingestion. We propose that honeybees may be behaviorally and physiologically adapted to tolerate low ethanol levels. Our findings highlight the need to consider both evolutionary adaptations and methodological sensitivity when evaluating affect-like states in insects. More sensitive continuous PER measures, explicit assays of sucrose responsiveness and motivation, and complementary neural or physiological markers will be essential to determine whether low-dose ethanol modulates arousal or affect-like processing in bees in other ways than tested here.

## 7. Compliance with Ethical Standards

No approval of research ethics committees was required to accomplish the goals of this study, because experimental work was conducted with an unregulated invertebrate species.

## 8. Supplementary Information

**Supplementary Information 1**. Diagram for exclusion/inclusion rationale.

**Supplementary Information 2**. Tables with additional results.

