## Supplementary figures and images for "Buzzed but not elated? Effect of ethanol on cognitive judgement bias in honeybees"

### Supplemental Information 1

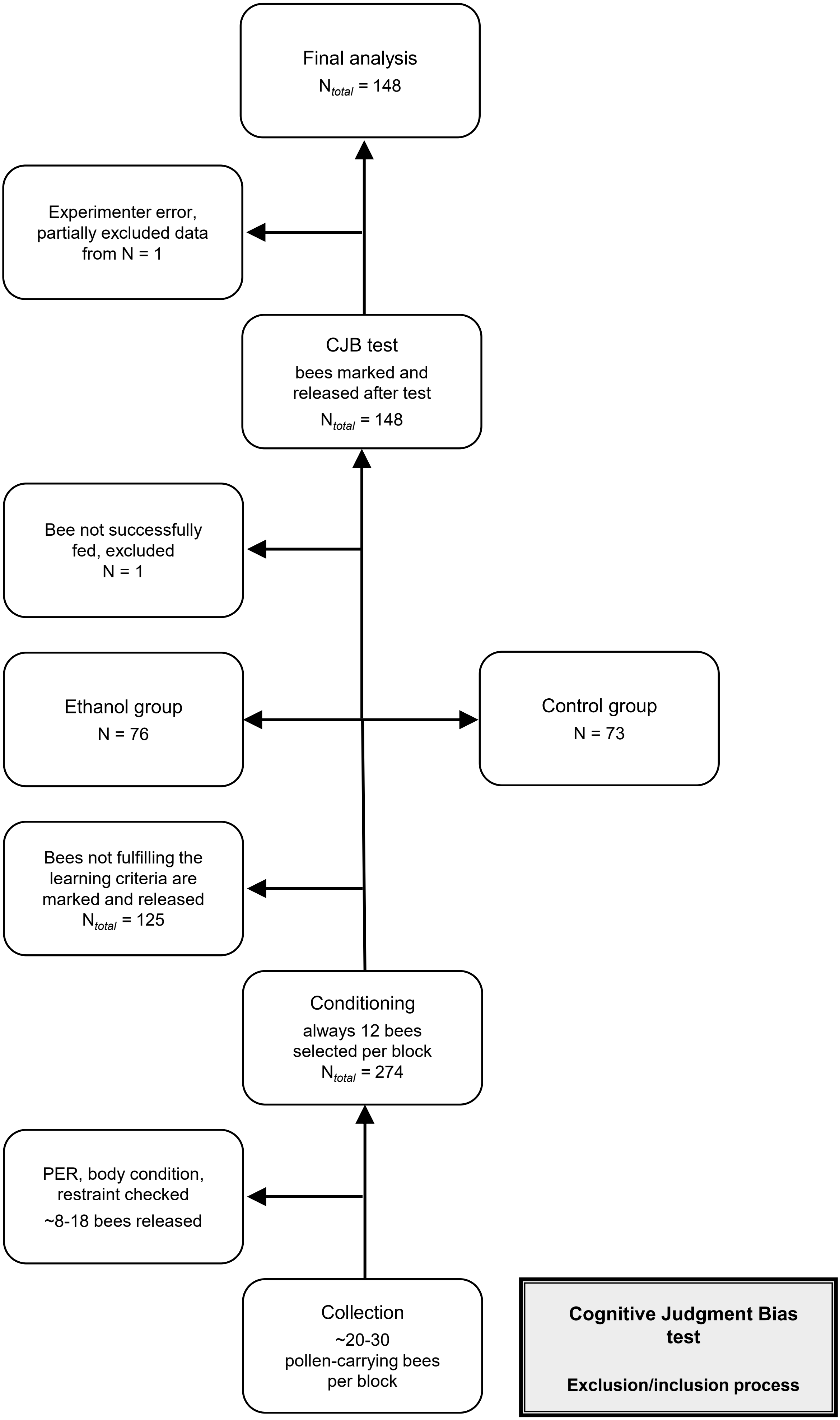

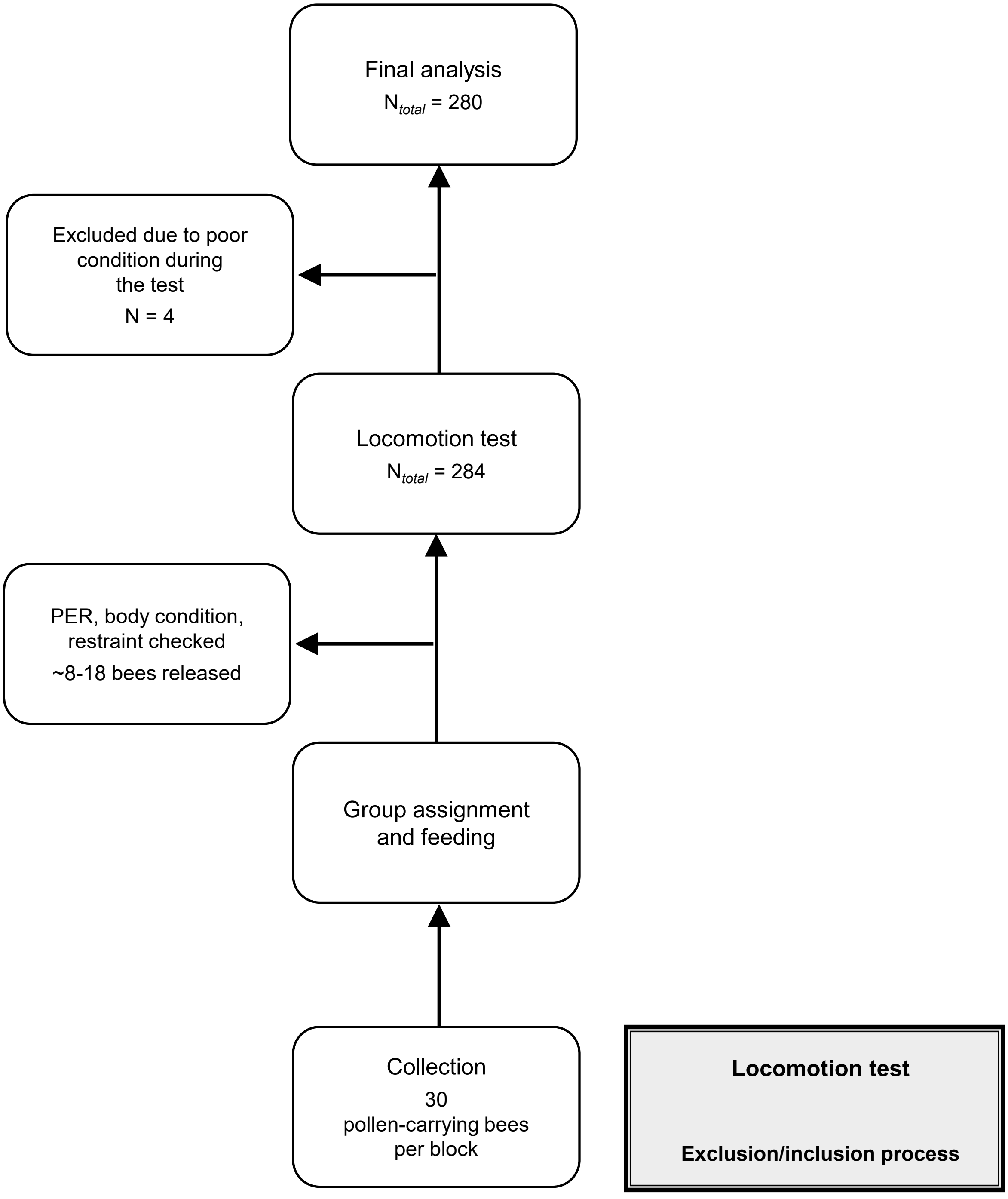

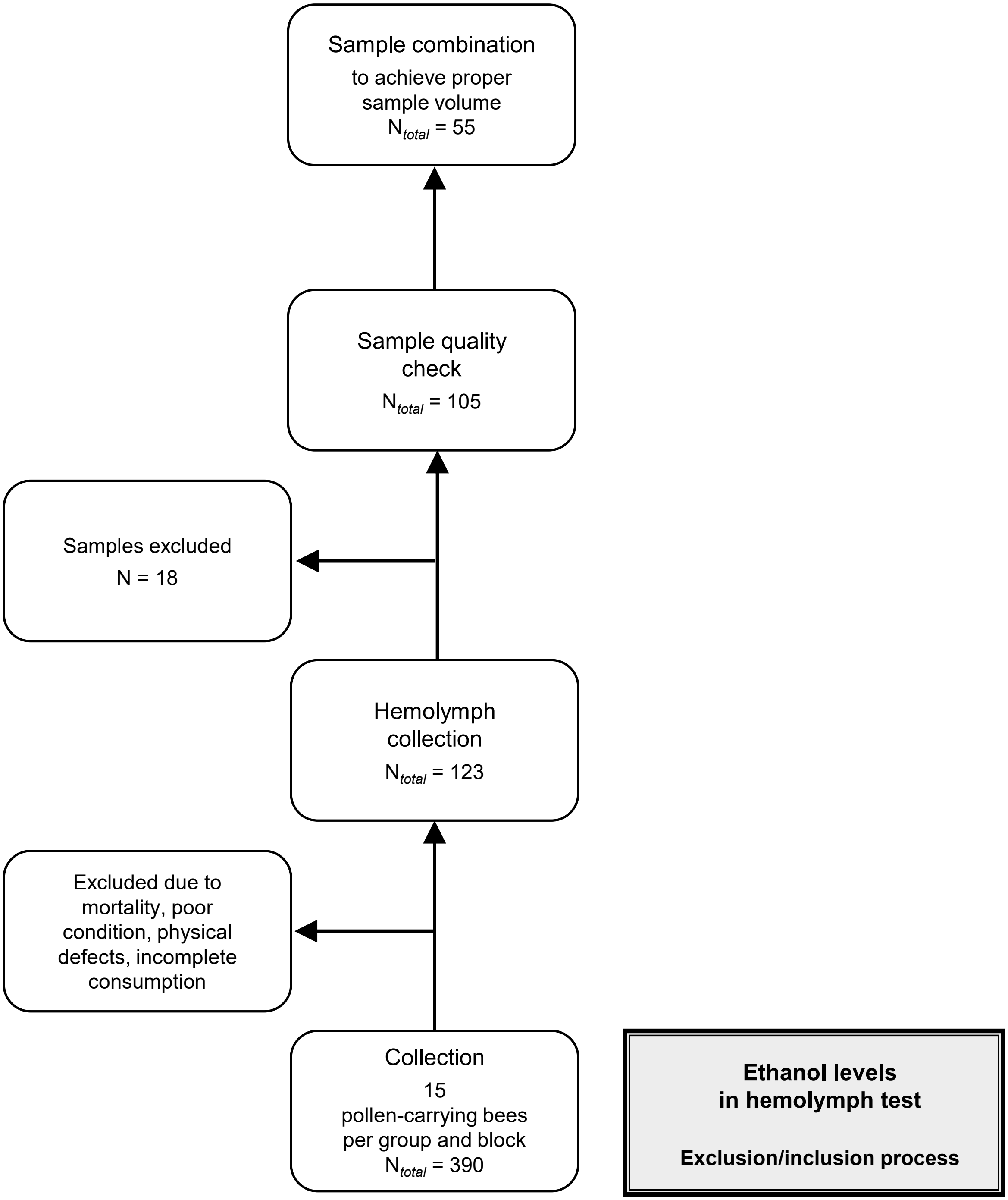
