## Supplemental Information 2 for "Buzzed but not elated? Effect of ethanol on cognitive judgement bias in honeybees"

Supplementary Material 2

**Table S1**. Contrasts in PER probability between the control and ethanol-fed bees at each odor mixture, based on the GAMM model of CJB. Formula: PER ~ group + s(odor_num, by = group, k = 4) + s(individual, bs = "re"). The column "mix" indicates consecutive tested odor mixtures (proportions of the appetitively conditioned odor octanone).

SE – Standard Error, CL – Confidence Limits (95% confidence level), OR – Odds Ratio with confidence intervals (95%)

| Mix | Estimate | SE | CL low | CL high | t | p | OR | OR low | OR high |
|---|---|---|---|---|---|---|---|---|---|
| 0 | -1.5802 | 1.1094 | -3.7585 | 0.5982 | -1.4244 | 0.1548 | 0.2059 | 0.0233 | 1.8188 |
| 2 | -0.1056 | 0.6039 | -1.2914 | 1.0803 | -0.1748 | 0.8613 | 0.8998 | 0.2749 | 2.9454 |
| 3 | -0.0241 | 0.4627 | -0.9326 | 0.8844 | -0.0520 | 0.9585 | 0.9762 | 0.3935 | 2.4216 |
| 4 | -0.0940 | 0.4459 | -0.9695 | 0.7815 | -0.2109 | 0.8330 | 0.9103 | 0.3793 | 2.1846 |
| 6 | 0.4282 | 0.4153 | -0.3872 | 1.2436 | 1.0311 | 0.3029 | 1.5345 | 0.6789 | 3.4682 |

**Table S2**. Analysis of Variance Table for the hemolymph ethanol model. Formula: log(1+EtOH) ~ group + family + plate + pool size.

df – degrees of freedom, SS – Sum of Squares, MS – Mean Square

| | df | SS | MS | F value | p |
|---|---|---|---|---|---|
| Group | 4 | 3.06822 | 0.76706 | 13.5852 | 2.432e-07 |
| Family | 3 | 0.33562 | 0.11187 | 1.9814 | 0.1303 |
| Plate | 1 | 0.10136 | 0.10136 | 1.7952 | 0.1870 |
| Individuals | 1 | 0.10549 | 0.10549 | 1.8683 | 0.1785 |
| Residuals | 45 | 2.54083 | 0.05646 | | |

**Table S3**. Full pairwise Tukey-adjusted contrasts between the control and ethanol-fed honeybees from the hemolymph ethanol model.

SE – Standard Error, CL – Confidence Limits (95% confidence level)

| Contrast | Estimate | SE | CL low | CL high | t | p |
|---|---|---|---|---|---|---|
| C - T120 | -0.399265 | 0.106824 | -0.70280 | -0.09572 | -3.73758 | 0.004528 |
| C - T240 | -0.303452 | 0.117270 | -0.63667 | 0.02976 | -2.58763 | 0.08984 |
| C - T30 | -0.652956 | 0.093568 | -0.91882 | -0.38708 | -6.97839 | 1.08e-07 |
| C - T60 | -0.491236 | 0.093832 | -0.75785 | -0.22461 | -5.23524 | 3.99e-05 |
| T120 - T240 | 0.095812 | 0.130559 | -0.27516 | 0.46679 | 0.73386 | 0.947406 |
| T120 - T30 | -0.253691 | 0.106047 | -0.55502 | 0.04763 | -2.39223 | 0.136176 |
| T120 - T60 | -0.091971 | 0.109208 | -0.40228 | 0.21833 | -0.84216 | 0.915906 |
| T240 - T30 | -0.349503 | 0.119904 | -0.69020 | -0.00880 | -2.91485 | 0.041833 |
| T240 - T60 | -0.187783 | 0.120513 | -0.53021 | 0.15464 | -1.55819 | 0.531242 |
| T30 - T60 | 0.161719 | 0.095374 | -0.10928 | 0.43272 | 1.69562 | 0.446855 |

**Table S4**. Probabilities, SEs, and CIs for the learning acquisition model. Formula: PER ~ CS_type * order + (1 | individual).

SE – Standard Error, Asymp. CL – Asymptotically estimated Confidence Limits (95% confidence level)

| CS_type | Order | Probability | SE | Asymp. CL low | Asymp. CL high |
|---|---|---|---|---|---|
| $CS^+$ | 1 | 0.000372 | 0.000303 | 0.000075 | 0.001836 |
| $CS^-$ | 1 | 0.003800 | 0.001626 | 0.001642 | 0.008773 |
| $CS^+$ | 2 | 0.028676 | 0.008853 | 0.015588 | 0.052172 |
| $CS^-$ | 2 | 0.001660 | 0.000857 | 0.000603 | 0.004562 |
| $CS^+$ | 3 | 0.232140 | 0.048552 | 0.150571 | 0.340198 |
| $CS^-$ | 3 | 0.000833 | 0.000522 | 0.000244 | 0.002839 |
| $CS^+$ | 4 | 0.368785 | 0.062620 | 0.256417 | 0.497453 |
| $CS^-$ | 4 | 0.000173 | 0.000189 | 0.000020 | 0.001475 |
| $CS^+$ | 5 | 0.373050 | 0.063180 | 0.259493 | 0.502574 |
| $CS^-$ | 5 | 0.000373 | 0.000305 | 0.000075 | 0.001851 |
| $CS^+$ | 6 | 0.526619 | 0.067325 | 0.395861 | 0.653826 |
| $CS^-$ | 6 | 0.000868 | 0.000543 | 0.000254 | 0.002957 |
| $CS^+$ | 7 | 0.741634 | 0.053393 | 0.624416 | 0.832106 |
| $CS^-$ | 7 | 0.000617 | 0.000430 | 0.000157 | 0.002418 |
| $CS^+$ | 8 | 0.703506 | 0.057831 | 0.579480 | 0.803365 |
| $CS^-$ | 8 | 0.000387 | 0.000316 | 0.000078 | 0.001921 |

**Table S5**. Analysis of robustness of the main model to the changes in the coding of outcome variable. Both models' formula: PER ~ group + s(odor_num, by = group, k = 4) + s(individual, bs = "re").

| | PER coded as: full = 1, partial or no response = 0 | PER coded as: full or partial response = 1, no response = 0 |
|---|---|---|
| **Smooth terms** | | |
| Control smooth | $\chi^2 = 65.53$, $p < 0.001$ | $\chi^2 = 65.37$, $p < 0.001$ |
| Group (ethanol) smooth | $\chi^2 = 84.95$, $p < 0.001$ | $\chi^2 = 83.95$, $p < 0.001$ |
| Individual smooth | $\chi^2 = 100.80$, $p < 0.001$ | $\chi^2 = 189.57$, $p < 0.001$ |
| **Parametric coefficients** | | |
| Intercept | $\beta = -2.26$, SE = 0.27, $z = -8.44$, $p < 0.001$ | $\beta = -1.33$, SE = 0.27, $z = -5.04$, $p < 0.001$ |
| Group (ethanol) | $\beta = -2.28$, SE = 0.42, $z = -0.66$, $p = 0.511$ | $\beta = 0.10$, SE = 0.37, $z = 0.27$, $p = 0.784$ |
| **Model fit** | | |
| Deviance explained | 51.6% | 50.9% |
| Adjusted $R^2$ | 0.503 | 0.508 |

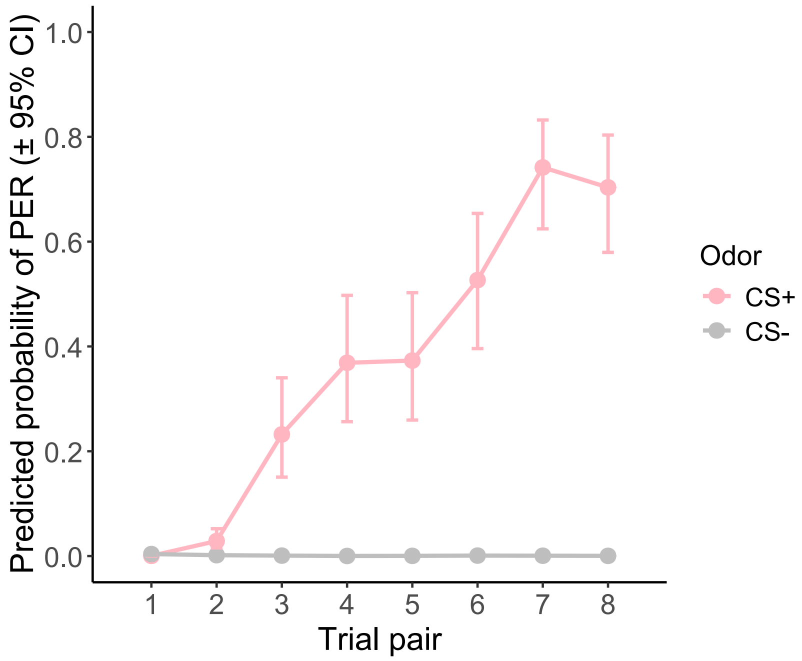


**Figure S1.** Acquisition of differential olfactory conditioning. Predicted probability of proboscis extension response (PER) to the rewarded ($CS^+$) and punished ($CS^-$) odors across eight conditioning trial pairs. Points and error bars represent model-estimated marginal means ± 95% confidence intervals derived from a binomial generalized linear mixed model with individual identity included as a random effect. PER responses to $CS^+$ increased sharply across trials, while responses to $CS^-$ remained low, demonstrating robust acquisition and discrimination learning.
